# Effect of partial selfing and polygenic selection on establishment in a new habitat

**DOI:** 10.1101/582122

**Authors:** Himani Sachdeva

## Abstract

This paper analyzes how partial selfing in a large source population influences its ability to colonize a new habitat via the introduction of a few founder individuals. Founders experience inbreeding depression due to partially recessive deleterious alleles as well as maladaptation to the new environment due to selection on a large number of additive loci. I first introduce a simplified version of the Inbreeding History Model (Kelly, 2007) in order to characterize mutation-selection balance in a large, partially selfing source population under selection involving multiple non-identical loci. I then use individual-based simulations to study the eco-evolutionary dynamics of founders establishing in the new habitat under a model of hard selection. The study explores how selfing rate shapes establishment probabilities of founders via effects on both inbreeding depression and adaptability to the new environment, and also distinguishes the effects of selfing on the initial fitness of founders from its effects on the long-term adaptive response of the populations they found. A high rate of (but not complete) selfing is found to aid establishment over a wide range of parameters, even in the absence of mate limitation. The sensitivity of the results to assumptions about the nature of polygenic selection are discussed.

## 1 Introduction

Peripheral habitats such as islands and geographic range limits present demographic and adaptive challenges to the establishment of new populations (Kawecki, 2008). Natural habitats often span environmental gradients, resulting in different selection pressures at the core and peripheries of the habitat. Peripheral habitats may also be subject to asymmetric gene flow, resulting in swamping, maladaptation and the emergence of ‘demographic sinks’ (Bridle and Vines, 2006). Alternatively, habitats colonized by a single long-distance dispersal event may be effectively isolated from the core, in which case the establishing population is strongly influenced by founder effects and prone to stochastic extinction. Other challenges stem from the low population densities that characterize initial phases of establishment. These result in increased inbreeding and associated fitness costs, while also rendering the population vulnerable to mate limitation and demographic Allee effects (Courchamp et al, 1999).

Several empirical studies have suggested a causal link between the mating system of a population and its establishment success in a new habitat. In a highly influential paper, Baker (1955) hypothesized that self-fertilizing species should be more adept at long-distance colonisation, and presented evidence for the over-representation of selfers on islands in comparison to the mainland. Subsequent work has revealed other examples of this general pattern (Barrett, 1996; Grossenbacher et al, 2017), but also important exceptions, notably the abundance of dioecious plants on the Hawaiian archipelago (Carlquist, 1966).

Arguments linking selfing to colonizing ability are primarily based on reduced mate limitation in selfing populations (Baker, 1955). Selfing, or more generally uniparental reproduction, provides *reproductive assurance*, allowing colonizers to survive the initial low-density phase (Pannell et al, 2015). However, mating systems affect several other aspects of establishment— complete or partial selfing changes the average heterozygosity along the genome, the extent of linkage and identity disequilibrium between loci under selection, and the amount of genetic variation in the population. These characteristics of the source population influence its adaptive potential in a new habitat, as well as the degree of inbreeding depression it might experience during the establishment bottleneck. Further, mating systems modulate outbreeding depression in the establishing population in the face of recurrent, maladaptive gene flow from the core habitat and thus, have the potential to themselves evolve under selection during establishment.

Given the many and possibly conflicting effects of mating system on establishment, theoretical models can play a crucial role in clarifying the range of environmental conditions and genetic parameters for which mating strategies such as increased selfing augment establishment success (Glémin and Ronfort, 2013; Uecker, 2017). An important challenge is to integrate polygenic architectures that often underlie adaptation into eco-evolutionary models that consider how population size and genotypic frequencies co-evolve.

Most theoretical work on the effects of polygenic adaptation during range expansions or the colonisation of new habitats has focused on *randomly mating* populations (Kirkpatrick and Barton, 1997; Polechova and Barton, 2015; Tufto, 2001; Barton and Etheridge, 2018). These models give insight into whether and how interactions and associations between loci-generated either by selection or due to mixing of diverged populations-impact evolutionary dynamics during establishment.

However, selfing and other forms of non-random mating also generate strong multi-locus associations. These have two major effects on a population under selection. First, correlations between homozygosity at different loci cause most deleterious alleles to be masked from selection in outcrossing or weakly selfing populations, but efficiently purged at higher selfing rates. Thus, allele frequencies and inbreeding depression exhibit a non-linear dependence on the selfing rate, especially when deleterious alleles are nearly recessive and the total mutation rate is high (Lande and Schemske, 1985; Lande et al, 1994). Second, selfing reduces homozygosity and the within-family variance of quantitative traits, while increasing their between-family variance (Wright, 1951). While the precise effect of selfing on quantitative trait variation depends on the magnitude and type of selection on the trait (Charlesworth and Charlesworth, 1995; Kelly, 1999; Lande and Porcher, 2015), adaptive response from quantitative variation is expected to be generally different in selfed versus outcrossed populations.

In this paper, I investigate how selfing within a large source population (e.g., on a mainland) influences its ability to colonize a new habitat (such as an island) in a scenario where the establishing population experiences both inbreeding depression and maladaptation to the new habitat due to selection on a large number of loci. For simplicity, it is assumed that environmental adaptation and inbreeding depression are affected by two *different* sets of unlinked loci. Alleles at the first set of loci have partially recessive effects and are unconditionally deleterious on both the mainland and the island. Alleles at the second set of loci have co-dominant effects and additively determine a trait which is under environment-dependent selection. The environmental trait is assumed to be under directional selection on both the mainland and island, but in opposite directions. The implications of these assumptions are explored in detail in the Discussion.

The study has two parts: I first use a simplified version of the inbreeding history model (Kelly, 2007) to characterize mutation-selection balance involving non-identical, unlinked loci under multiplicative selection in a large, partially selfing source population. The focus is on elucidating the extent to which associations between loci are explained by differences in *recent* selfed versus outcrossed ancestry of individuals.

In the second part, I explore how the genetic composition of a large source population influences establishment probabilities on the island, following the introduction of a few founder individuals from the source. Successful establishment requires that the population both survive increased inbreeding depression (due to higher levels of inbreeding in small populations, and segregation of partially recessive alleles) and adapt (via a response from existing genetic variation or new mutations). The goal is to understand how selfing within the source population affects both these aspects of the establishment process, and explain the resulting dependence of establishment probabilities on selfing rate. Another goal is to distinguish the effect of selfing on the *initial fitness* of founders from its effect on how variable or inbred their descendants are, which determines the long-term *adaptive potential* of the population.

The interplay between partial selfing and polygenic selection in large populations has been analysed using different theoretical approaches (Kondrashov, 1985; Charlesworth et al, 1990, 1991; Lande et al, 1994; Kelly, 1999, 2007; Roze, 2015; Lande and Porcher, 2015; Awad and Roze, 2018). The main challenge is to find tractable and accurate approximations for the multi-locus associations that emerge due to partial selfing even in the absence of linkage. Roze (2015) and Awad and Roze (2018) derive analytical expressions for allele frequencies under different kinds of selection by assuming that these are only affected by *pairwise* associations between loci. This analysis is thus applicable for low genome-wide mutation rates, nearly co-dominant loci, and weak selfing, but becomes inaccurate if these conditions are not met (see fig. 4 in Roze (2015)), which generates significant multi-locus disequilibria between loci.

An interesting approach by Kelly (1999, 2007) classifies individuals according to their *selfing age*, i.e., the number of generations of continuous selfing in the lineage leading up to the individual. The partially selfing population can then be viewed as a structured population consisting of groups or cohorts of individuals of different selfing ages. Kelly (2007) used this approach to analyse a model with identical loci subject to partially recessive, deleterious mutations. He derived recursions for the mean and variance of (and the correlation between) the number of loci that are homozygous and heterozygous for the deleterious allele within each selfing age cohort by assuming that associations, i.e., linkage and identity disequilibria *within* cohorts are weak. The underlying assumption is that in the absence of linkage and epistasis, variation of inbreeding coefficients between individuals in the population is mostly due to differences in their recent selfing histories.

The present work employs a simpler approximation which neglects disequilibria within cohorts altogether, but accounts for disequilibria that emerge across the whole population due to differences in average allele frequencies or average homozygosity between cohorts. This approximation is thus slightly less accurate than that of Kelly (2007), but has the advantage of yielding simpler recursions which can be easily generalized to describe the evolution of *non-identical* loci. As shown below, ignoring associations within cohorts yields reasonably accurate predictions for allele frequencies, pairwise associations between loci, and mean fitness and inbreeding depression in the population across a range of parameters. This also allows us to predict the genetic composition of source populations with different selfing fractions, without directly simulating large numbers of individuals with many selected loci.

While the effects of inbreeding *during* establishment have been studied in recent theoretical work (Barton and Etheridge, 2018), the implications of having systematic deviations from panmixia in the source population itself remain largely unexplored. Dornier et al (2008) consider how inbreeding depression and Allee effects shape the establishment potential of partially selfing populations by assuming a fixed level of inbreeding depression. However, as demonstrated below, establishment success depends on the interplay between inbreeding depression and the fitness of founders, which are correlated in a complex way when the total genomic mutation rate is high. Moreover, establishment often involves adaptation to a new environment via response from quantitative genetic variation. The consideration of source populations with complex genetic architectures and non-random mating is thus an important step towards modeling more realistic population establishment or evolutionary rescue scenarios.

Since the main goal is to understand how selfing affects establishment probability via the genetic composition of founding individuals subject to polygenic selection, we will only consider a single bout of migration. We thus ignore the effect of selfing on outbreeding depression as well as heterosis, which may, however, be important when the establishing population is subject to continuous gene flow from a divergent source population. Further, the analysis will focus on *initial* establishment: this distinction is important, since selfing may have different effects in small and growing versus large and equilibrated populations. Finally, selfing rates on the island are assumed to be the same as in the source population. Thus, the model does not allow for mating system plasticity or the evolution of selfing rates in the new habitat, which could, however, be important during the establishment of natural populations (Peterson and Kay, 2014).

## Model and Methods

### Source population

Consider a large, partially selfing source population with *N* diploid, hermaphroditic individuals. Each individual genome has *L*_*A*_ loci (referred to as additive loci henceforth) which undergo mutation between two alternative alleles with *co-dominant* effects, and *L*_*R*_ loci (referred to as recessive loci) which undergo mutation to deleterious alleles with *partially recessive* effects. The co-dominant alleles contribute additively to a trait *z* under directional selection. All loci are unlinked, and there is no epistasis between loci. Mutation between the two alternative allelic states occurs at rates *μ*_*A*_ and *μ*_*R*_ per locus per generation for the additive and partially recessive loci respectively.

For simplicity, effect sizes of the two alternative alleles are set to −*α*/2 and *α*/2 at each additive locus (though the approximations and results generalize to the case where effect sizes are unequal across loci). The trait value *z* thus ranges from *z*_*min*_= −*αL*_*A*_ to *z*_*max*_ = *αL*_*A*_. The effect *α* can be arbitrarily set to 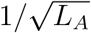 in accordance with the usual quantitative genetics convention. For simplicity, it is assumed that all deleterious recessive alleles are also characterised by the same selective disadvantage *s* and dominance coefficient *h*, with *h* < 1/2. Individual fitness is then given by 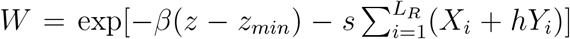 where *X*_*i*_ and *Y*_*i*_ are equal to 1 respectively if the individual is homozygous or heterozygous for the recessive allele at locus *i*, and zero otherwise. The strength of selection per allele at each additive locus is thus *s̃* = *βα*. It is sometimes convenient to use the negative log fitness *G* = −ln(*W*), which is the genetic load associated with an individual, and which is the sum of two components: *β*(*z* − *z*_*min*_) and 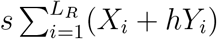.

Generations are assumed to be non-overlapping. The lifecycle in each generation consists of mutation, followed by selection, and then mating via partial self-fertilization (in which a fraction *r*_*s*_ of individuals self). Since fitness is *multiplicative* across both types of loci, and loci are unlinked, there should be no multi-locus associations in a sufficiently large population that is either purely outcrossing (*r*_*s*_ = 0) or purely selfing (*r*_*s*_ = 1). However, partial selfing (0 < *r*_*s*_ < 1) generates associations between allelic states (linkage disequilibrium or LD) as well as between homozygosity (identity disequilibrium or ID) at different loci even in the absence of epistasis, linkage and drift (Weir and Cockerham, 1973).

### Identity and Linkage Equilibrium within Cohorts (ILEC) approximation

Such associations arise due to differences in selfing histories and the resultant variation in homozygosity across individuals in a partially selfing population. Following Kelly (2007), we can define the *selfing age* of an individual as the number of generations back to its most recent outcrossing ancestor, or equivalently, the number of generations of continuous selfing in the lineage leading up to the individual. Thus, an individual produced by outcrossing in the present generation has selfing age 0, the selfed offspring of a parent produced via outcrossing in the previous generation has selfing age 1, and so on. Individuals with higher selfing ages have higher homozygosity on average.

Let the proportion of individuals with selfing age *i* be *f*_*i*_, and the average frequency of homozygous loci (with two ‘1’ alleles) among these individuals be 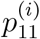. Then, under the assumption that the states of different loci are uncorrelated within cohorts, the identity disequilibrium between a pair of loci (of the same type) across the whole population is given by (Supporting Information):

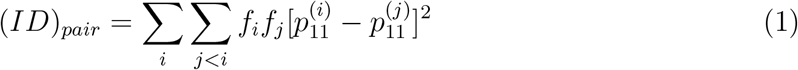

Here, the double summation is over all possible pairs of selfing ages. Thus, population-wide disequilibria arise due to the presence of cohorts with different average homozygosities and allele frequencies per locus, even when there are no associations within cohorts.

In general, each such cohort is itself characterized by some population structure— for instance, the cohort with selfing age zero consists of outcrossed offspring of parents with diverse selfing histories (and hence slightly different allele frequencies), which generates LD within this cohort. However, in the present approximation, all multi-locus associations (LD and ID) within a cohort are neglected. Then the state of the population is completely specified by the fraction *f*_*i*_ of individuals belonging to cohort *i*, the frequencies of homozygous and heterozygous additive loci (denoted by 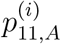 and 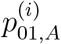 respectively) within the *i*^*th*^ cohort, and the corresponding frequencies 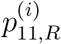 and 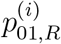 for partially recessive loci. This will be referred to this as the Identity and Linkage Equilibrium within Cohorts (ILEC) approximation, to distinguish it from the inbreeding history model (IHM) introduced by Kelly (2007). Note that the latter also accounts for weak, pairwise disequilibria within cohorts.

Under the ILEC approximation, the evolution of the partially selfing population is described by specifying how the proportions *f*_*i*_, and the frequencies 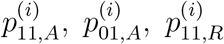 and 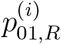 change due to mutation, selection and partial selfing in each generation (see SI). These *deterministic* equations ignore allele frequency changes due to drift, as well as stochastic fluctuations of the proportions *f*_*i*_, and are thus only applicable for large source populations. These equations can be iterated over generations, until equilibrium is attained. The equilibrium frequencies within each cohort and the corresponding fractions *f*_*i*_ can then be used to calculate all population-wide disequilibria (e.g., eq. (1), see also SI), as well as the full fitness distribution in the population, under the ILEC assumption.

### Individual-based simulations

The key assumption underlying the ILEC approximation is that a single round of outcrossing is sufficient to erase most associations between loci, and that the residual associations can be ignored for prediction of population attributes. This assumption is tested by simulating large populations for various parameter combinations.

Simulations are initialized by choosing the allelic state of each locus for each of the *N* individuals independently. The population is evolved in discrete generations as follows— first, all individuals undergo mutation, where the allelic state of each locus is flipped (0 ↔ 1) with probability *μ*_*R*_ for a recessive locus and probability *μ*_*A*_ for an additive locus. *N* individuals are then chosen for mating by sampling (with replacement) from the population with probabilities proportional to individual fitness. Each individual is then allowed to self with probability *r*_*s*_ or outcross with probability 1 −*r*_*s*_. For outcrossing individuals, the mating partner is chosen by again sampling individuals in proportion to their fitness. All parental individuals produce gametes via free recombination of their diploid genomes. Selfed offspring are then created by pairing gametes from the same individual and outcrossed offspring by pairing gametes from the two (different) parental individuals.

The population is evolved for a few thousand generations until there is no further change in allele frequencies and disequilibria. For each set of parameters, reliable estimates of various quantities of interest are obtained by averaging over several replicates. All statistics are measured at the end of the generation. Comparisons with individual-based simulations show that the ILEC approximation predicts detailed attributes of the source population such as pairwise disequilibria between loci, as well as the *distribution* of genetic load among individuals in the population with reasonable accuracy (figs. 1 and 2 below).

**Figure 1:**
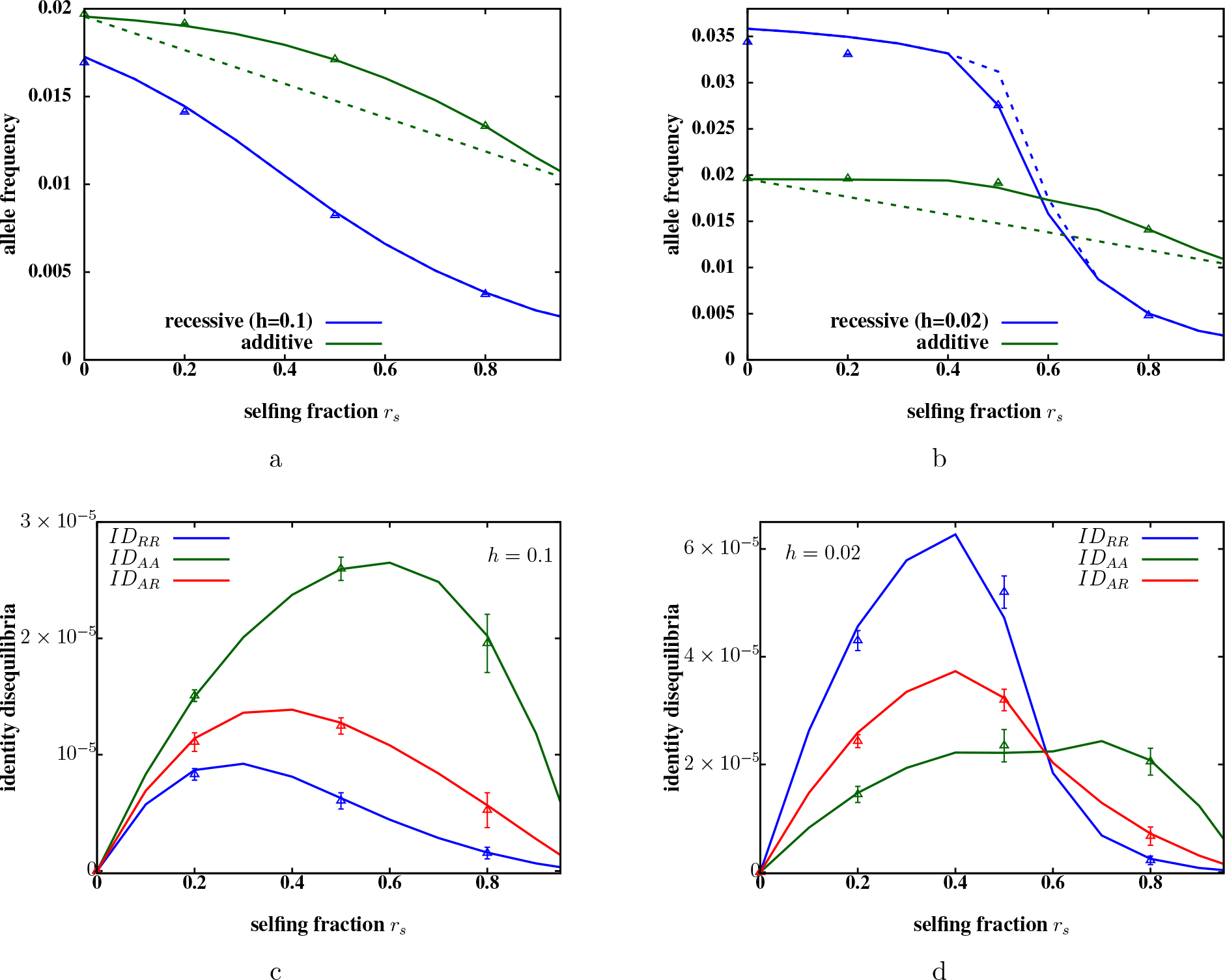
(A)-(B) Equilibrium allele frequencies at partially recessive (blue) and additive (green) loci versus selfing fraction *r_s_*, when the genome has *L*_*A*_= 1000 additive and *L*_*R*_= 5000 partially recessive loci. The dominance coefficient of partially recessive alleles is *h* = 0.1 in fig. A, and *h* = 0.02 in fig. B. All loci are unlinked. Predictions of the ILEC approximation (solid lines) agree closely with results from simulations of *N* = 10, 000 individuals (triangles). Dashed lines are the corresponding allele frequencies (obtained from the ILEC approximation) when only one type of loci is present— thus, the green dashed line represents additive allele frequencies in a genome with 1000 additive loci (and no recessive loci). Presence of unlinked deleterious recessive mutations inflates the frequency of the unfavourable additive allele (dashed vs. solid green lines), especially for intermediate selfing fractions. However, the frequency of recessive alleles is not strongly affected by unlinked additive alleles (dashed and solid blue lines are indistinguishable in fig. A). (C)-(D) Various pairwise identity disequilibria versus *r*_*s*_ for *h* = 0.1 (fig. C) and *h* = 0.02 (fig. D) respectively. Green, red and blue solid lines show the ILEC predictions for ID between two additive loci on the genome (*ID*_*AA*_), or two recessive loci on the genome (*ID*_*RR*_), or between an additive and a recessive locus (*ID*_*AR*_); triangles show the corresponding disequilibria, as obtained from individual-based simulations of a population with *N* = 10000. The mutation rate per locus is *μ*_*A*_ = *μ*_*R*_= 10^−4^, selection against partially recessive deleterious alleles is *s* = 0.05 and against additive alleles is *s̃* = *βα* = 0.005.

### Population establishment in the new habitat

In the second part of the paper, I investigate how founders from source populations with different selfing fractions colonize a new environment. Since establishment typically involves a few individuals and proceeds via a phase of small population size, we cannot use the deterministic ILEC approximation and must explicitly account for drift and demographic stochasticity by simulating individuals.

However, founders from a large source population can still be drawn using the ILEC approximation: each founder is assigned a selfing age *i* with probability equal to the proportion *f*_*i*_ of individuals belonging to cohort *i* in the population. The proportions {*f*_*j*_} depend on selection, dominance and mutation parameters, and the selfing rate in the source population, and can be obtained from the ILEC approximation, as described above (see also SI). Then, each of the *L*_*A*_ additive loci in the founder genome is independently assigned one of three possible genotypes : 00, 01/10 or 11 (corresponding to two possible alleles) with probabilities 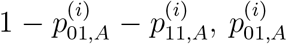 or 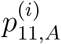 respectively. Here, 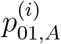 and 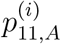 denote the frequency of additive loci that are heterozygous and homozygous for the ‘1’ allele, within the *i*^*th*^ cohort. Recessive loci are assigned genotypes similarly, i.e., based on the equilibrium heterozygote and homozygote frequencies 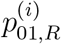 and 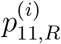 in cohort *i*. Choosing genotypes independently at each locus reflects the simplifying assumption that there are no associations between loci within cohorts of individuals with a given selfing age.

Establishment is initiated by a single founder event in which *N*_0_ individuals from the source population are introduced all at once into the new habitat. There is no subsequent migration. The direction of selection on the additive trait is reversed in the new habitat (with respect to the source population), such that individual fitness in the new habitat is: 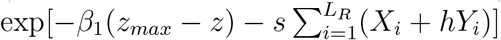, where *β*_1_ is positive, and is typically different from *β*_0_, the strength of selection in the source population. Contrast this with the fitness function in the source population: while partially recessive alleles are unconditionally deleterious in both the source population and the new habitat, different additive alleles are favoured in the new habitat versus the source population. Thus, additive alleles have *environment-dependent* fitness effects. The establishing population is subject to *hard* selection in the new habitat, such that mean fitness influences population size.

Establishment in the new habitat is studied via individual-based simulations. These are initialized by randomly sampling *N*_0_ founder genomes from the source population, as described above. Mutation is implemented as before. Hard selection is enforced by assuming that the total number *N*_*t*+1_ of offspring produced in generation *t* + 1 is a Poisson-distributed random variable with mean given by 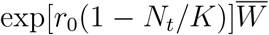. Here, *r*_0_ is the intrinsic rate of growth of the population, *N*_*t*_ is the number of individuals prior to selection, *K* is the carrying capacity of the new habitat, and 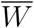 is the mean population fitness, obtained by averaging over the fitness of all *N*_*t*_ individuals in the new habitat.

Each of the *N*_*t*+1_ offspring is assumed to be produced via selfing (with probability *r*_*s*_) or outcrossing (probability 1−*r*_*s*_). One (or two) parent(s) of each selfed (or outcrossed) offspring are chosen from among the *N*_*t*_ individuals by sampling with probabilities proportional to fitness. Note that if *N* (*t*) is small, then the same individual may be drawn both times while sampling the two parents of an outcrossed offspring. Thus, the realised selfing fraction may be much higher than *r*_*s*_— being 1 if there is a single individual in the parental generation, and approaching *r*_*s*_ as population size increases. Gametes are generated via free recombination, and paired to produce the next generation of individuals, as in the source population.

To assess the colonisation potential of a source population, a thousand independent colonisation events are simulated. Each event involves *N*_0_ founders, independently sampled from the source. Establishment probability in the new habitat is then computed as the fraction of ‘successful’ establishment events among these. Establishment is considered successful if the population size is at least *K*/10 individuals at the end of a certain time period (here taken to be 100 generations) after the founder event. Since density-dependence has little effect in such small populations (*N/K* ~ 0.1), these simulations yield approximately the same results as a simpler model where each individual has a Poisson-distributed number of offspring with mean equal to its fitness multiplied by the growth rate *r*_0_ (Barton and Etheridge, 2018).

## Results

### Mutation-selection balance in the source population: ILEC approximation

We will first analyze attributes of a large source population (neglecting drift) under partial selfing and polygenic selection. Figures 1a and 1b show the equilibrium frequencies of the negatively selected allele at the two types of loci, in an example where fitness is affected by both. Allele frequencies obtained from simulations of 10000 individuals (points) are in close agreement with predictions of the ILEC approximation for both *h* = 0.1 and *h* = 0.02, across a range of selfing fractions.

The high rate of recessive mutations (*U*_*R*_ = 2*μ*_*R*_*L*_*R*_ = 1) relative to the (weak) selective effect per allele (*U*_*R*_/*hs* equal to 200 and 1000 in figs. 1a and 1b respectively), results in the segregation of a large number of recessive alleles. This generates substantial fitness differences between selfed and outcrossed individuals within a population, especially in weakly selfing populations (see also fig. 2d). Thus selfers tend to be significantly under-represented (relative to the selfing fraction *r*_*s*_) among parents of the next generation of offspring. This implies that most deleterious alleles are masked from selection, since selection against deleterious alleles is less effective within the outcrossing cohort as compared to selfing cohorts, especially for low *r*_*s*_ and when the average heterozygosity is high. As a consequence of this kind of *selective interference* between alleles, deleterious alleles are purged efficiently only at high selfing fractions, when selfed individuals make a non-negligible genetic contribution to the next generation of offspring (Lande et al, 1994). The ILEC approximation captures the threshold selfing fraction, beyond which purging is effective, with reasonable accuracy (fig. 1b), unlike calculations that only include corrections due to pairwise disequilibria (Roze, 2015).

**Figure 2:**
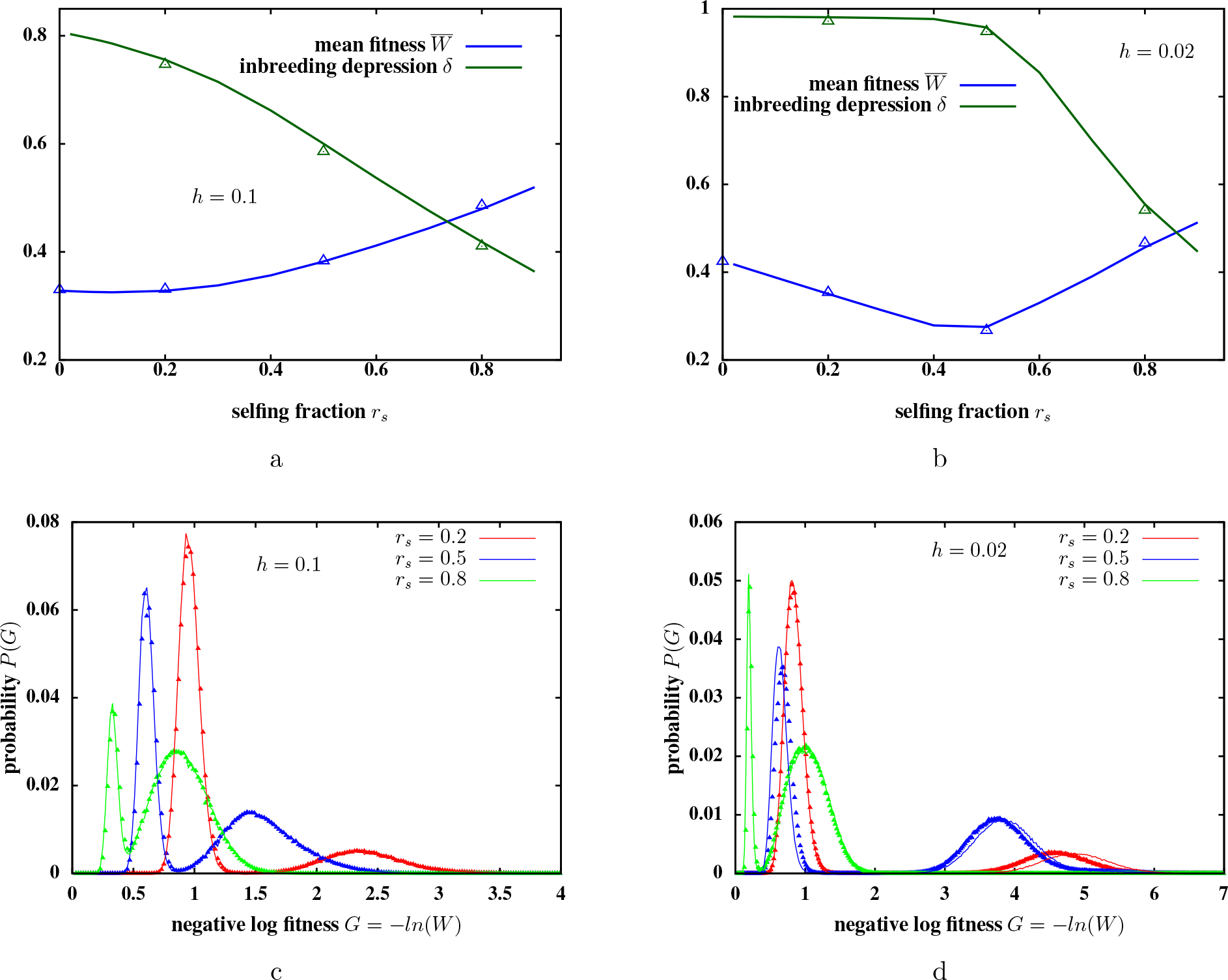
(A)-(B) Mean population fitness 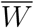 and inbreeding depression *δ* versus selfing fraction *r_s_*, when the genome has *L*_*A*_ = 1000 additive and *L*_*R*_= 5000 partially recessive loci. The dominance coefficient of partially recessive alleles is *h* = 0.1 in fig. A, and *h* = 0.02 in fig. B. Solid lines represent predictions of the ILEC approximation, triangles depict results from simulations of *N* = 10000 individuals. (C)-(D) Comparison of simulation results (triangles) and ILEC predictions (lines) for the probability distributions of genetic load *G* (defined as negative log fitness *G* = − ln(*W*)) in the source population. The plots show load distributions for three different selfing fractions: *r*_*s*_= 0.2, 0.5, 0.8, and for two different dominance values of the recessive allele: *h* = 0.1 (fig. C) and *h* = 0.02 (fig. D). The distribution of *G* is bimodal due to significantly higher number of homozygous deleterious, recessive alleles in the genomes of selfed versus outcrossed offspring. Mutation rates per locus are *μ*_*A*_ = *μ*_*R*_ = 10^−4^, selection against partially recessive deleterious alleles is *s* = 0.05 and the strength of directional selection is *s̃* = *βα* = 0.005.

A related effect is observed at additive loci, where the frequency of unfavourable alleles is *inflated* by unlinked deleterious recessives segregating in the population (in figs. 1a and 1b, compare solid lines which represent allele frequencies in a genome having both additive and recessive loci with dashed lines which represent allele frequencies in a genome with only one type of locus). This is again a consequence of high inbreeding depression due to recessive alleles, which strongly reduces the contribution of selfed individuals to the gametic pool from which the next generation of offspring is formed. As a result, the effective selection against unfavourable additive alleles is weaker than it would be in the absence of recessive alleles. This effect is typically quite modest, and is most significant at intermediate *r*_*s*_, for which the mean and variance of the additive trait may increase by as much as 20 – 25% due to unlinked deleterious recessive alleles. More generally, unlinked recessive alleles affect additive trait variation in a complex way that depends qualitatively on *U*_*R*_(to be explored in detail elsewhere).

The ILEC approximation is inaccurate for *r*_*s*_ close to 1: purely selfing populations tend to fix deleterious alleles due to smaller effective population sizes *N*_*e*_ (Charlesworth et al, 1993). In particular, *N*_*e*_ is strongly reduced close to *r*_*s*_ = 1, when the total genomic mutation rate is high relative to selection, i.e., *U*_*R*_/*s* ≫ 1. This builds up negative disequilibria between deleterious alleles, which decreases fitness variance and the efficacy of selection in the population (Kamran-Disfani and Agrawal, 2014). This effect is not captured by the deterministic ILEC approximation, which does not account for LD due to drift. Thus, equilibrium allele frequencies are several times higher than the ILEC prediction for *r*_*s*_∼ 1 and *U*_*R*_/*s* = 20 (not shown in figs. 1a and 1b), even for population sizes as large as 10000.

The ILEC approximation also predicts identity and linkage disequilibria between alleles at different loci. Figures 1c and 1d compare pairwise ID obtained from simulations with the corresponding ILEC prediction, and show that the approximation is accurate for ID between two loci of the same type (blue and green triangles), as well as between loci of different types (red triangles). As expected, ID is strongest for intermediate selfing fractions when the distribution of selfing ages and inbreeding coefficients in the population is widest, resulting in maximally structured populations. Pairwise linkage disequilibria are found to be negligible for the same parameters (except for *r*_*s*_ ∼ 1) both in simulations and according to the ILEC prediction.

The ILEC approximation can also predict the average population fitness 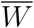 and the inbreeding depression *δ*, defined as 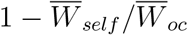 (where 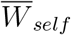 or 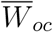 is the average fitness of a randomly chosen selfed or outcrossed individual). Note that 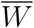 and *δ* depend on the full genotypic distribution, and are thus affected by all disequilibria (at least when selective interference between loci is strong). As before, there is good agreement between simulation results (triangles) and ILEC predictions (lines) for various selfing fractions and dominance values (figs. 2a and 2b).

For *h* = 0.02, the mean fitness is minimum at intermediate selfing fractions (fig. 2b). An increase in *r*_*s*_ reduces the frequency of deleterious alleles (which tends to increase fitness), while increasing the average homozygosity (which tends to reduce fitness). Since highly recessive alleles are effectively masked from selection at low selfing fractions in the selective interference (*U*_*R*_/*hs* ≫ 1) regime, the reduction in deleterious allele frequency with *r*_*s*_ is quite modest (see fig. 1b). Thus, increased selfing reduces fitness at low *r*_*s*_ primarily by generating excess homozygosity. The ineffectiveness of selection at low *r*_*s*_ is also reflected in the fact that inbreeding depression only starts falling beyond a threshold *r*_*s*_ (fig. 2b).

Further, we can use the ILEC approximation to generate the *distribution* of load in the population (see SI for details) and compare this with equilibrium distributions from individual-based simulations (figs. 2c and 2d). Here, load is simply negative log fitness 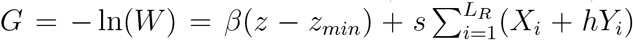 and is the sum of two components, the first due to additive alleles that influence environmental adaptation, and the second due to unconditionally deleterious recessive mutations. The ILEC prediction is very accurate for higher dominance (*h* = 0.1) but slightly less so when alleles are more recessive (*h* = 0.02).

A key feature of the load distribution is that it is bimodal, with outcrossed individuals having significantly lower load due to recessive alleles than individuals with one or more generation(s) of continuous selfing in their lineage. This difference is especially marked when alleles are highly recessive and selfing fractions small or intermediate. Note that cohorts with different selfing have different average homozygosity. However, these differences are comparable to the variance of homozygosity within each cohort, so that the fitness distributions of cohorts with different *non-zero* selfing ages overlap significantly, resulting in a single broad peak at high load (figs. 2c and 2d).

### Population establishment in the new habitat

To understand how selfing influences establishment in the new habitat, it is useful to consider scenarios where genetic load is either only due to unconditionally deleterious recessive alleles or only due to locally maladaptive additive alleles, and then analyse a scenario with selection on both. We will investigate establishment for a range of selfing fractions from *r*_*s*_ = 0 to *r*_*s*_= 0.9, but will not consider complete selfing (*r*_*s*_ = 1), as this leads to high fixation rates of deleterious mutations, even in large source populations (Kamran-Disfani and Agrawal, 2014).

### Establishment scenario with environment-independent selection on recessive alleles

For simplicity, deleterious alleles at each locus are assumed to have the same selective effect *s* and dominance value *h*, in each source population. Moreover, *s* and *h* are environment-independent, i.e., are the same in the new habitat. Thus, in this scenario, establishment does not involve adaptation to a new environmental optimum, but only requires that the establishing population purge the excess genetic load that arises from increased inbreeding just after colonisation.

Since the *N*_0_ founder genomes are generated using the deterministic ILEC approximation, there is no identity by descent in the population at *t* = 0. I further consider only those parameters *U_R_*, *s* and *h* for which a large source population would be viable under hard selection (with the same intrinsic growth rate *r*_0_). This is ensured by testing that for each parameter combination, a population with *N*_0_ = 100 individuals doubles with probability greater than 0.95 within 100 generations, for very large *K*. In principle, *N*_0_ = 100 is small enough that drift, stochastic fluctuations and inbreeding may be significant. Thus, this is a rather conservative criterion for testing the viability of a *large* population.

Figure 3a shows how the establishment probability *P*_*est*_ varies with the selfing fraction *r*_*s*_ of the source population in an example with *N*_0_ = 10 founders. Note how the dependence of *P*_*est*_ on *r*_*s*_ changes qualitatively with the recessivity and selective effect of deleterious alleles. When genetic load is due to nearly recessive, weakly selected deleterious alleles, then *P*_*est*_ is minimum for intermediate *r_s_*. By contrast, when populations undergo mutation to less recessive or more strongly deleterious alleles, then *P*_*est*_ increases monotonically with *r_s_* (neglecting the *r_s_ ~* 1 behaviour).

**Figure 3:**
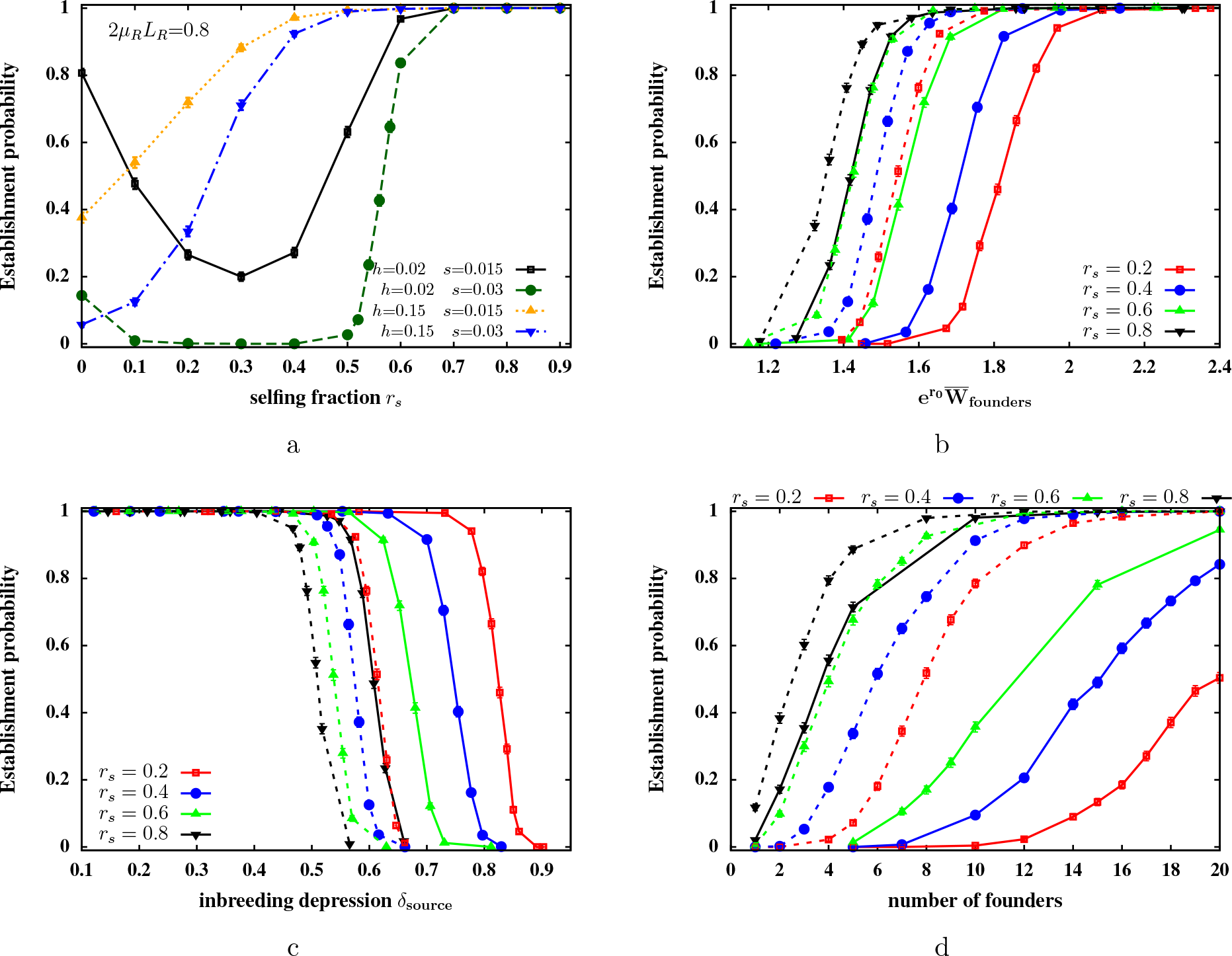
Establishment probabilities *P*_*est*_ in the new habitat in a scenario where founders only carry deleterious recessive alleles with environment-independent selective effects. (A) *P*_*est*_ versus selfing fraction *r*_*s*_ for different selective effects *s* and dominance values *h* of deleterious mutations. *P*_*est*_ is minimum at intermediate *r*_*s*_ if genetic load is due to weakly selected, nearly recessive alleles, but increases monotonically with *r*_*s*_ for larger *h*. Simulation parameters: *L*_*R*_ = 4000, *μ*_*R*_ = 10^−4^. (B)-(C) *P*_*est*_ as a function of the initial growth rate of founders (fig. B) and as a function of inbreeding depression in the source population (fig. C), for different selfing fractions (represented by different colors) and different dominance coefficients (solid lines for *h* = 0.02 versus dashed lines for *h* = 0.1). The initial growth rate of the founders and inbreeding depression in the source are tuned by changing the total mutation rate *U_R_*. The number of founders is *N*_0_ = 10. (D) *P*_*est*_ versus the number of founders *N*_0_ for source populations with different *r*_*s*_ and different dominance values (solid and dashed lines for *h* = 0.02 and *h* = 0.1 respectively). The mutation rate *UR* is chosen independently for each source population such that all populations have the same mean fitness 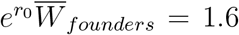. Simulation parameters in figs. (B)-(D): *L*_*R*_ = 4000, *s* = 0.02. *P*_*est*_ is the fraction of successful establishment events among 1000 independent simulation runs, each initialized by generating *N*_0_ founders from the source population using the ILEC approximation. Growth rate is *r*_0_ = 1.1 in the new habitat; carrying capacity is *K* = 1000.

Since selection pressures on the mainland and island are identical, and we have only considered parameter combinations for which a large source population would be viable under hard selection, failure to establish in the new habitat must arise solely from inbreeding depression due to the small number of founders, and cannot be due to their low initial fitness. However, the extent to which inbreeding depression reduces establishment probability does depend on the initial fitness of founders— even moderate inbreeding depression can prevent establishment if the initial founder fitness is close to the threshold of viability 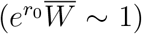, while very fit founders 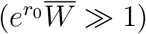 would establish despite high inbreeding depression.

Thus, the complex dependence of *P*_*est*_ on selfing fraction reflects the underlying dependence of both the initial fitness of founders and the magnitude of inbreeding depression on *r_s_*. For highly recessive alleles and high genomic mutation rates *U_R_*, fitness is minimum at intermediate selfing fractions in a large population (fig. 2b). Moreover, inbreeding depression shows little dependence on *r*_*s*_ for weak selfing. Thus, founders with intermediate *r*_*s*_ are least fit and experience similar levels of inbreeding depression as founders with *r*_*s*_ = 0, which explains the minimum in *P*_*est*_ at intermediate *r*_*s*_ for *h* = 0.02 (fig. 3a). For less recessive alleles or lower mutation rates *U_R_*, the mean fitness of a large partially selfing population increases and the inbreeding depression decreases with *r*_*s*_(fig. 2b). Thus, outcrossing populations are at maximum disadvantage, resulting in a monotonic increase in *P*_*est*_ with *r*_*s*_ for *h* = 0.15 in fig. 3a.

To further disentangle the effects of selfing fraction on founder fitness and inbreeding depression, we can plot *P*_*est*_ as a function of the initial growth rate of the founders (fig. 3b). The initial growth rate is 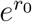 multiplied by the mean fitness of the founders in the new habitat, which, in this scenario, is just their mean fitness in the source population. Figure 3b shows that *P*_*est*_ becomes non-zero above a threshold founder fitness which depends on both *r*_*s*_ and the dominance coefficient *h*. For a given *h*, the threshold fitness required for establishment decreases with *r_s_*. This just reflects the fact that a strongly selfing population with the same mean fitness as a weakly selfing population, must have lower heterozygosity, and hence, experience a lower level of inbreeding depression. Similarly, for a given selfing fraction, the threshold founder fitness is lower when mutations are less recessive (solid versus dashed lines), due to the lower inbreeding depression associated with higher values of *h*. The dependence on *h* is especially marked in weakly selfing populations.

It is also informative to plot *P*_*est*_ as a function of inbreeding depression *δ*_*source*_ in the source population (fig. 3c). As expected, establishment is successful only below a threshold value of *δ_source_*. Figure 3c further shows that this threshold for inbreeding depression is actually *lower* for populations with higher selfing fractions. This is due to the fact that a strongly selfing population must harbour more deleterious alleles on average and thus have lower fitness than a weakly selfing source population with the same level of inbreeding depression. In addition, populations with the same same selfing fraction *r*_*s*_ and the same level of inbreeding depression have establishment probabilities which depend on the recessivity of alleles contributing to inbreeding depression: *P*_*est*_ is higher if alleles are *more* recessive (dashed vs. solid lines).

Since the transient increase in inbreeding just after colonisation depends crucially on the number *N*_0_ of founders, it is useful to consider how *P*_*est*_ varies with *N*_0_, for founders drawn from source populations which have the same mean fitness but different selfing fractions (fig. 3d). Consistent with 3b, high rates of selfing allow for establishment with fewer founders, because of the lower inbreeding depression *δ*_*source*_ associated with these founders. Further, *P*_*est*_ increases more slowly with *N*_0_ for more recessive alleles, again because of higher *δ*_*source*_ associated with smaller values of *h*.

### Establishment scenario with environment-dependent selection on additive alleles

Consider now a scenario where individuals only carry loci that mutate between alternative co-dominant alleles that additively determine a trait *z*. The fitness of an individual with trait value *z* is proportional to exp[−*β*_0_(*z − z_min_*)] in the source population, and exp[−*β*_1_(*z_max_ − z*)] in the new habitat, where *β*_0_ and *β*_1_ are both positive. Thus, in this scenario, establishment is primarily constrained by maladaptation of the founders to the new environment, and is aided by the ability of the population to adapt from standing variation. Figure 4a shows how *P*_*est*_ varies with the selfing fraction of the source population, following a single colonisation event involving *N*_0_ = 10 founders.

In the absence of deleterious recessive mutations, the frequency of the locally unfavourable additive allele is approximately ~ (*μ_A_/*2*s̃*_0_)(2 − *r_s_*) in the source population (Ohta and Cockerham (1974), see also dashed lines in figures 1a and 1b). As a result, stronger selfing reduces the frequency of alleles that are selected against in the source and conversely, favoured in the new habitat (where the direction of selection on the additive trait is reversed). Thus, other parameters being the same, founder fitness in the new environment declines with *r_s_*, which causes establishment probabilities to also decline with *r*_*s*_ (fig. 4a). This dependence on selfing fraction arises only close to a threshold value of *s̃*_1_ = *β*_1_*α*, for which the genetic load of founders in the new habitat, given by 2*s̃*_1_*L_A_*[1 − (*μ_A_/*2*s̃*_0_)(2 − *r_s_*)], is comparable to the growth rate *r*_0_. Populations fail to establish, irrespective of selfing fraction, when selection in the new habitat is very strong (2*s̃*_1_*L_A_ ≫ r*_0_). Conversely, for 2*s̃*_1_*L_A_ ≪ r*_0_, founders from any source population have high establishment success.

As before, we can ask: does selfing influence establishment probability predominantly through the initial fitness of founders or more via its effect on the rate of adaptation of the establishing population? Figure 4b shows *P*_*est*_ as a function of initial fitness of founders, which is varied by varying the mutation target *U*_*A*_ = 2*μ*_*A*_*L*_*A*_ for the additive trait. For a given number of founders *N*_0_, the curves for different values of *r*_*s*_ appear to collapse onto a single curve, suggesting that in this situation, establishment probabilities depend on the selfing fraction only via its effect on the fitness of the founders. Contrast this with the previous scenario with deleterious recessive mutations, where the threshold fitness required for establishment exhibited a marked dependence on *r*_*s*_(fig. 3b).

This is surprising at first glance since it suggests that adaptation plays little role during establishment. To investigate this further, we can follow the dynamics of the average population size 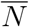 and the average genetic load 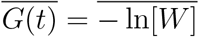 (figures 4c and 4d) for four different groups of founders with different *r*_*s*_ but the same mean fitness (here 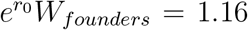). Note that the genetic load declines (or equivalently, the frequency of the favoured alleles increases) faster in the more strongly selfing populations, from the outset (fig. 4c). However, this does not affect the *initial* growth rate of the population (fig. 4d). More precisely, if the initial growth rate of founders *r*_0_ − *G*_0_ is high compared to the rate of adaptation −*dG*(*t*)*/dt*, populations rapidly attain a size ~ *K*(1 − *G*(0)*/r*_0_), more or less independently of the actual rate of adaptation. As a result, initial establishment (which is defined here as reaching a minimum size of ~ *K/*10) depends only on the initial fitness of the founders in fig. 4d (where (1 − *G*(0)*/r*_0_) ~ 0.135). However, *long-term* population growth does depend on the rate of adaptation and is thus faster in populations with higher selfing fractions.

**Figure 4:**
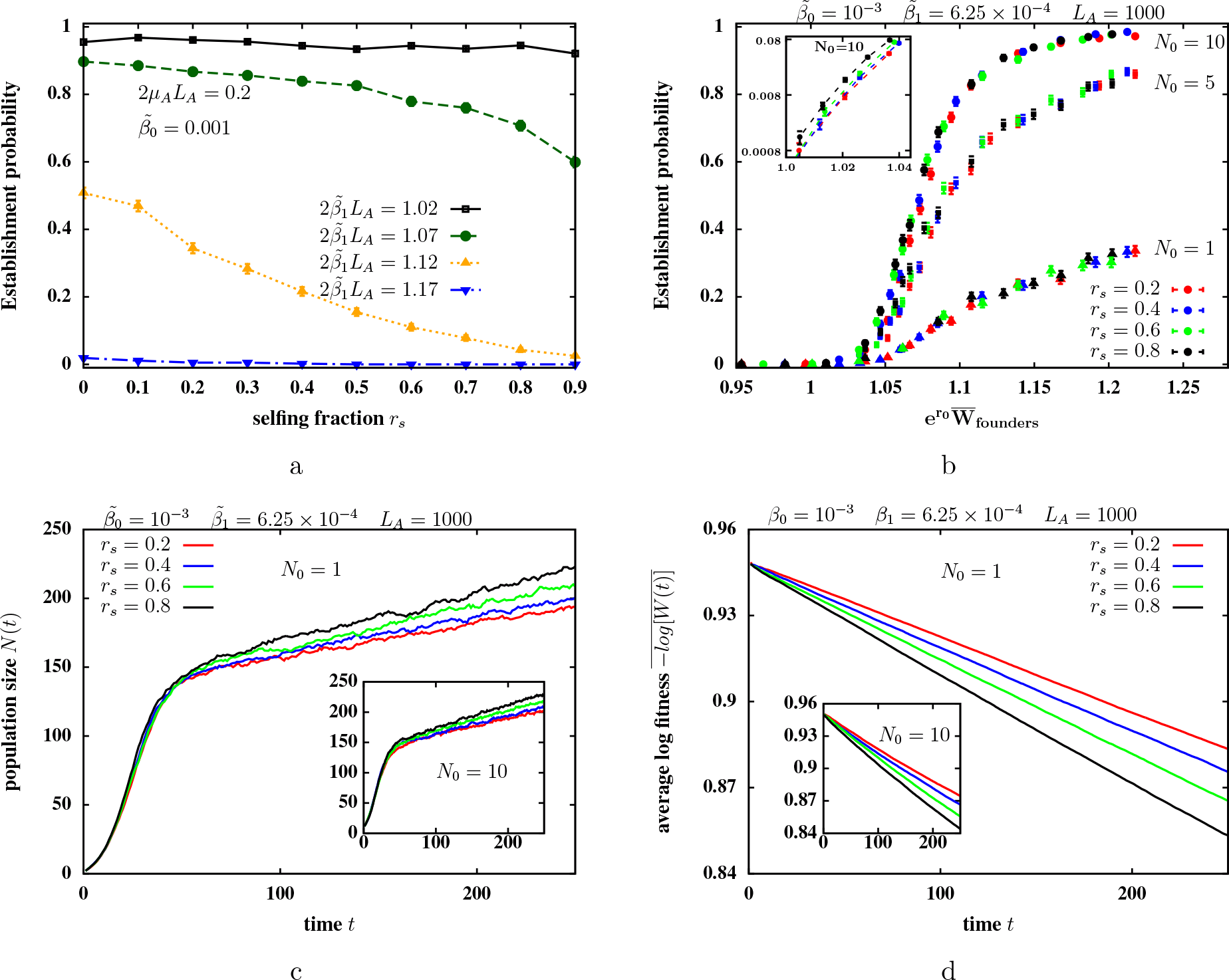
Establishment probabilities *P*_*est*_ in the new habitat in a scenario where founders only carry co-dominant alleles that additively determine a trait under environment-dependent selection. (A) *P*_*est*_ versus selfing fraction *r*_*s*_ for different selection strengths (expressed as 2*s̃*_1_*L*_*A*_) in the new habitat with *N*_0_ = 10 founders. *P*_*est*_ declines with increasing *r*_*s*_ for intermediate 2*s̃*_1_*L*_*A*_. Simulation parameters: *L*_*A*_ = 1000, *μ*_*A*_ = 10^−4^, *s̃*_0_ = 0.001. (B) *P*_*est*_ as a function of the initial growth rate of founders in the new habitat for various *r*_*s*_ (represented by different colors) and various values of *N*_0_ (represented by different symbols). The initial growth rate of the founders is varied by changing the total mutation rate *U*_*A*_ = 2*μ*_*A*_*L*_*A*_, which changes the minor allele frequency. The inset zooms into the parameter region with rare establishment (*P*_*est*_ 1) for *N*_0_ = 10. (C)-(D) Average population size 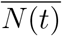 (fig. C) and average load or negative log fitness 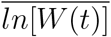 (fig. D) in the new habitat versus time *t*, when founders have the same mean fitness 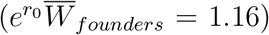 in the new habitat, but different selfing fractions *r*_*s*_ and genomic mutation rates *U*_*A*_. The quantities 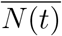 and 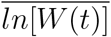 are calculated by averaging over those replicates in which the population is not extinct until 250 generations. Main plots and insets depict dynamics for *N*_0_ = 1 and *N*_0_ = 10 respectively. Genetic load decreases faster in highly selfing populations but population growth is insensitive to *r*_*s*_ at short time scales. Simulation parameters for figs. (B)-(D): *L*_*A*_= 1000, *β*_0_ = 10^−3^, *β*_1_ = 6.25 × 10^−4^. Intrinsic growth rate is *r*_0_ = 1.1 in the new habitat; carrying capacity is *K* = 1000.

The above reasoning suggests that when the initial fitness advantage of founders is comparable to the rate of adaptation, then selfing fraction should influence initial establishment more strongly. However, for weak selection per locus (which implies slow adaptation), this is precisely the condition under which establishment would be improbable. To test this, we zoom into the parameter region with 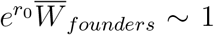 (see inset of fig. 4b), and find that selfing has a more significant effect on *P*_*est*_ when establishment is rare. For instance, in the inset of fig. 4b, the establishment probability of founders drawn from a source population with *r*_*s*_ = 0.8 is ~ 1.75 times that of founders with *r*_*s*_ = 0.2 that have the same fitness in the new habitat.

### Establishment scenario with both types of alleles

Finally consider establishment scenarios where founders carry both unconditionally deleterious, partially recessive alleles and additive alleles under environment-dependent selection. Figure 5 compares source populations with different total mutation rates *U_R_*, and hence different magnitudes of genetic load due to partially recessive variants. For each *U_R_*, we can find the critical environmental selection strength *β*_1*,c*_, such that *P*_*est*_ is significant (greater than 0.05), as long as the selection strength in the new habitat is weaker than *β*_1*,c*_ (see fig. 5a). A high value of *β*_1*,c*_ signifies that the population can establish despite a large reversal of environmental selection.

As expected, for any selfing fraction, the range of environmental selection strengths, to which a population can adapt, shrinks as *U*_*R*_ increases. When *U*_*R*_ is close to zero, outcrossing populations (*r_s_* ~ 0) are able to adapt to slightly larger shifts than highly selfing populations (*r*_*s*_ = 0.8). However, for any other small, non-zero value of *U_R_*, founders from the *r*_*s*_ = 0.8 population establish over a larger range of *β*_1_. This is due to the lower deleterious recessive load and lower inbreeding depression in large, highly selfing populations. Note that this is true for almost completely recessive alleles (*h* = 0.02, solid lines in fig. 5a) as well as partially recessive alleles (*h* = 0.2, dotted lines).

We can also measure different components of the genetic load (negative log fitness − ln(*W*)) associated with founders, for parameter combinations with *P_est_* > 0.05. The genetic load is the sum of two components— one arising from deleterious recessive mutations, and the other from local maladaptation of the additive trait. The first component has average value 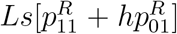 (where 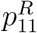 and 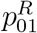 are the homozygote and heterozygote frequencies of the recessive allele in the source), and the second component has average value 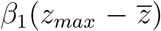 (where 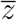 is the population average of the additive trait in the source). Figure 5b depicts the values of these two components on a two dimensional phase plot: for instance, 10 founders drawn from a source population with *r*_*s*_= 0 and dominance *h* = 0.02 of the recessive alleles, establish a colony in the new habitat with probability greater than 0.05, only for points (representing the two load components) lying below the solid red line.

Figure 5b shows that the total load is a good predictor of establishment success when deleterious mutations are less recessive and source populations strongly selfing. However, for weakly selfing populations, which suffer significant inbreeding depression due to highly recessive alleles, establishment success cannot be predicted on the basis of the total genetic load of the founders, but requires a consideration of the different contributions to load. Thus, for low values of *r*_*s*_ and *h*, the threshold total fitness required for establishment is significantly higher (or maximum possible load significantly lower) when there is a high rate of deleterious recessive mutations.

**Figure 5:**
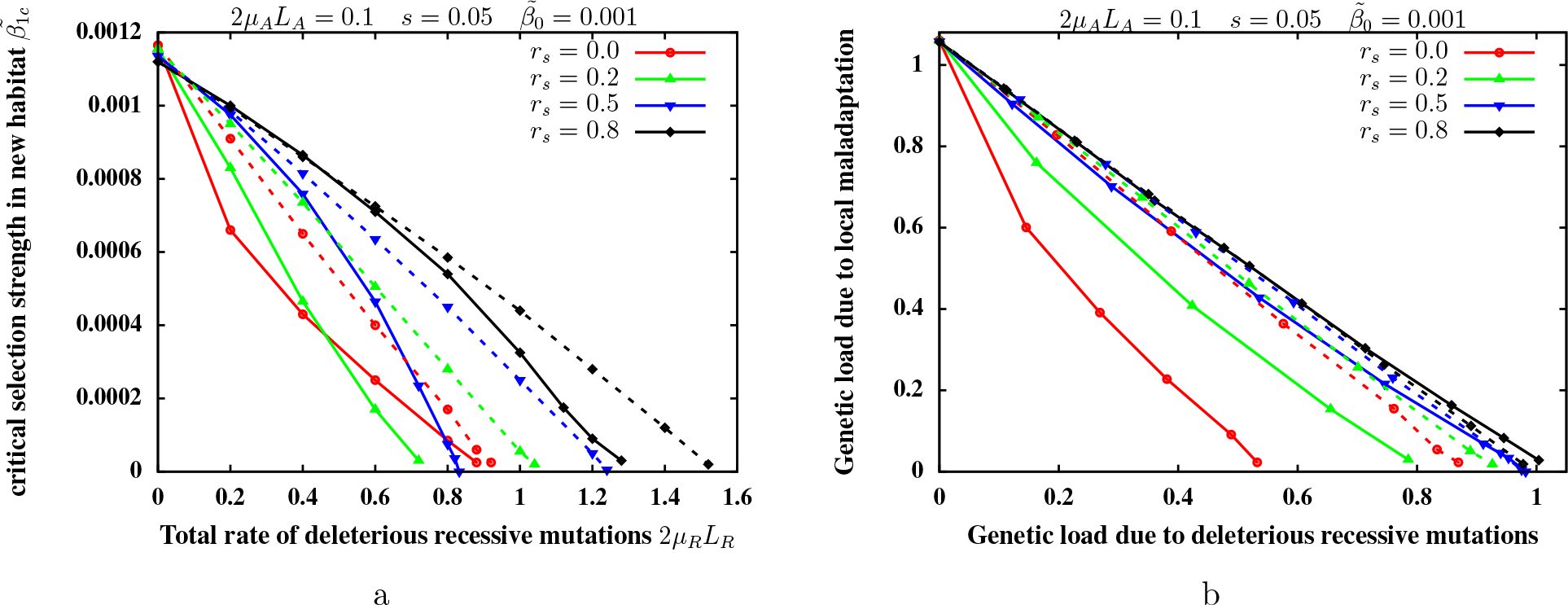
Phase diagrams showing parameter combinations for which the establishment probability *P*_*est*_ is greater than 0.05, when individual genomes contain both partially recessive alleles with environment-independent deleterious effects, and additive alleles with environment-dependent effects. (A) Critical selection strength *s̃*_1*,c*_ on the additive trait in the new habitat versus the total mutation rate *U*_*R*_ for the partially recessive alleles for different *r*_*s*_ and *h*. For a given *U_R_*, the establishment probability *P*_*est*_ is greater than 0.05 only if *s̃*_1_ < *s̃*_1*,c*_, i.e., in the parameter regions below the corresponding line. (B) The maximum (average) genetic load due to maladaptation in the new habitat for which establishment is possible (*P_est_* > 0.05), as a function of the genetic load due to deleterious recessive alleles. Solid and dashed lines depict phase boundaries when deleterious alleles have dominance values *h* = 0.02 and *h* = 0.2 respectively; different colors show phase boundaries for different values of *r_s_*. Establishment probabilities are obtained as the fraction of successful establishment events out of 1000 independent simulation runs, each initialized by sampling *N*_0_ = 10 founders from the source population. Growth rate is *r*_0_ = 1.1 in the new habitat; carrying capacity is *K* = 1000. Other parameters are: *L*_*A*_ = 500, *μ_A_* = 10^−4^, *L*_*R*_ = 2000, *s* = 0.05, *s̃*_0_ = 0.001.

## Discussion

Partial selfing is common across a variety of taxa, with some analyses reporting intermediate selfing rates (between 0.2 and 0.8) in as many as 40 − 50% of seed plants (Goodwillie et al, 2005) and hermaphroditic animal species (Jarne and Auld, 2006). The extent of inbreeding depression in populations with intermediate selfing rates may be similar to that in outcrossers, especially among long-lived taxa such as gymnosperms with high per-generation mutation rates (Winn et al, 2011). This suggests a highly polygenic architecture of inbreeding depression, characterized by selective interference between recessive alleles (Lande et al, 1994), in many natural populations. Extensive work on the genetics of inbreeding depression points towards an important role of both highly recessive lethals and moderately recessive, weakly deleterious alleles (Charlesworth and Willis, 2009). More generally, estimates in *Drosophila melanogaster* and *Saccharomyces cerevisiae* reveal a wide distribution of dominance values of deleterious alleles, with a mean of 0.1-0.2 (Agrawal and Whitlock, 2011; Peters et al, 2003), and also suggest a negative correlation between the dominance values of deleterious alleles and selection against them (Charlesworth, 1979).

The present study represents a preliminary attempt to understand how the interplay of selfing and polygenic architecture shapes the eco-evolutionary dynamics of a population establishing in a new habitat. Our model makes several assumptions about the genetic architecture of the establishing population, whose implications are examined below. First, the model assumes that inbreeding depression and response to environmental selection are due to distinct, non-overlapping sets of unlinked loci. Founders are drawn from source populations in mutation-selection balance, with directional selection on the environment-dependent trait. Further, source populations are assumed to be large enough that mutation-selection balance is essentially deterministic, with a negligible role for drift. Under these assumptions, high rates of (but not complete) selfing is found to facilitate establishment in several distinct ways.

### Establishment in the presence of deleterious recessive mutations

Large populations harbour a substantial number of deleterious recessive alleles when the genomic rate of deleterious mutations *U*_*R*_ is high with respect to the typical selective effect *hs*. In this scenario, colonies founded by a small number of individuals may fail due to a kind of genetic Allee effect, wherein an increase in the fraction of selfed individuals (over and above *r_s_*) or increased mating between related individuals depresses fitness, which reduces population size and further increases inbreeding, ultimately resulting in extinction. This effect is ameliorated if the source population is itself highly selfing— inbreeding depression declines significantly with *r*_*s*_ for *U_R_* ≫ *hs*, which gives highly selfing founders an advantage over equally fit, weakly selfing founders (fig. 3b).

More generally, our analysis points towards the difficulty of predicting establishment success based only on average fitness of the source population or on inbreeding depression alone. In a partially selfing population, these two quantities bear no simple relationship to each other, when selective interference between loci is strong, unlike in the case with *U_R_/hs* ~ 1 (Bataillon and Kirkpatrick, 2000). Founders with the same initial fitness show different degrees of inbreeding depression *δ* that depend on *s*, *h*, *r*_*s*_ and *U_R_*. Conversely, founders drawn from populations with the same *δ* have different mean fitness and thus are affected by inbreeding depression to different extents. In particular, the threshold value of inbreeding depression *δ_c_*, beyond which establishment becomes unlikely, is higher in outcrossing versus selfing populations with the same mean fitness (fig. 3c).

The success of highly selfing founders in establishing despite the initial bottleneck hinges on the effective purging of deleterious variants in large source populations with high *r_s_*. Purging is less effective, however, when the source population is itself small (Glémin, 2003), as is often the case in human-assisted re-introduction of endangered species into new habitats. Understanding how the genetic composition of *small* source populations influences their establishment potential remains an important challenge in conservation biology.

The realised rate of selfing or biparental inbreeding during initial establishment depends crucially on the effective number of founders. In the present model, this is equal to or less than 2*N*_0_– this could describe, for instance, the establishment of a diploid plant population via dispersal of seeds into a new habitat. In an alternative scenario, where populations are founded by *N*_0_ fertilized adults (carrying sperm from one or many fathers), the effective number of founders could be larger than 2*N*_0_. Importantly, in our model, founders are *not* obligate outcrossers; thus, founders from source populations with *r*_*s*_ = 0, can nevertheless self under conditions of mate limitation, resulting in a severe *genetic* Allee effect. If the source population is self-incompatible, then the realised rate of selfing in the new habitat is zero, irrespective of *N*_0_. In this case, outcrossers suffer much less from inbreeding depression during establishment, but are subject to a *demographic* Allee effect, wherein population growth rate is strictly zero for *N*_0_ = 1. This leads to an even stronger advantage for self-compatible founders for low *N*_0_.

### Establishment with adaptation from standing additive variation

Selfing has a qualitatively different effect when the establishing population must adapt to a different environment via a response from standing variation and new mutations at additive loci. We have analysed a specific scenario in which the direction of selection on the environmental trait is reversed in the new habitat, so that the response to selection occurs only via rare alleles. In this scenario, if the number of loci *L*_*A*_ is large, then selection per locus must be correspondingly weak for initial founder fitness to be in a range where establishment is possible. As a consequence, adaptation is too slow to significantly affect establishment, which is instead determined essentially by the initial fitness advantage of founders. Thus, the selfing fraction in the source population influences establishment probability only via founder fitness, except when *P*_*est*_ is small (fig. 4b).

In an alternative scenario, where the trait is under stabilizing selection towards different optima in the two habitats, adaptation to the new habitat would involve a response from both rare and common alleles. In this scenario, if the difference between the two selection optima is small relative to the standard deviation of trait values among founders, then adaptation can play a more important role during initial establishment than in the present model. As before, adaptation should be faster in more strongly selfing populations due to stronger effective selection per favourable allele. However, with strong stabilizing selection, co-dominant alleles also generate inbreeding depression: selfed cohorts have higher trait variance and lower fitness than outcrossed cohorts, and thus may make little genetic contribution to the next generations. As a result, trait variance is unaffected by selfing at low selfing rates but purged at higher selfing rates (Lande and Porcher, 2015). Understanding how selfing in a source population under stabilizing selection influences its colonisation potential via effects on genetic variation, effective population size and rate of adaptation is an interesting avenue for future work.

More generally, when the trait is determined by a large number of small-effect loci, the genotypic values of the offspring of any two individuals are approximately normally distributed (Barton et al., 2017). In the absence of selfing, this allows for a description of the eco-evolutionary dynamics of an establishing population within an infinitesimal framework (Barton and Etheridge, 2018), which is characterised by relatively few parameters such as the population size, the mean and variance of the trait under selection, and the distribution of inbreeding coefficients in the population. In principle, this framework can be extended to include selfing (although incorporating dominance within the infinitesimal model is more challenging). A key feature of selfing is that it *redistributes* variance (from within families to between families), unlike inbreeding due to small population sizes (as considered by Barton and Etheridge (2018)) which reduces both within and between family variance. Thus, the two forms of inbreeding are expected to have qualitatively different effects on establishment.

### Establishment involving both inbreeding depression and adaptation to a new environment

Under the assumptions of this model, highly selfing populations establish over a wider parameter region, especially when the total rate *U*_*R*_ of deleterious mutations is large, and mutations highly recessive. Importantly, with low or intermediate *r_s_*, establishment success depends not only on founder fitness, but also on what proportion of fitness loss is due to recessive alleles (fig. 5b).

The present model is relatively easy to analyse since it makes the convenient assumption that the alleles that contribute to local adaptation and the alleles that are unconditionally deleterious are distinct and unlinked. In reality, variants that affect fitness are often highly pleiotropic. Thus, an alternative model would be one where alleles at a large number of loci affect multiple traits under stabilizing selection. In this case, most alleles are deleterious and have recessive effects on fitness *on average*, though with a wide variance of dominance coefficients (Manna et al, 2011). A shift in the selection optima for one or more traits in the new habitat would then necessitate a response from variants with a range of adaptive effects and contributions to inbreeding depression. This model could thus provide an alternative framework for studying the interplay between inbreeding depression and polygenic adaptive response during establishment.

Linkage between adaptive and deleterious alleles could also qualitatively change our results. In particular, strong selfing significantly reduces the effective rate of recombination when loci are tightly linked. This generates Hill-Robertson interference between adaptive alleles, which reverses the advantage selfers experience during adaptation from co-dominant or even mildly recessive alleles (Hartfield and Glémin, 2016). Linkage also increases hitchhiking of deleterious variants with adaptive alleles, especially in highly selfing populations, which reduces the fixation probability of even slightly recessive adaptive alleles (Hartfield and Glémin, 2014; Kamran-Disfani and Agrawal, 2014). An interesting question is whether linkage could thus reduce the selfing fraction that is ‘optimal’ for establishment in new habitats.

Our analysis focuses on establishment that involves a single founder event. A natural extension is to study establishment via recurrent migration from the source population (Barton and Etheridge, 2018). Continual gene flow is expected to alleviate the high inbreeding depression experienced by predominantly outcrossing populations via heterosis, and thus reduce the advantage of highly selfing founders during initial establishment. On the other hand, a highly selfing population, once established, might better withstand maladaptive gene flow from the mainland and experience less outbreeding depression. An interesting question is whether different mating strategies might be favoured by natural selection during different phases of establishment.

For simplicity, the analysis included only two kinds of loci. However, the ILEC approximation provides a computationally frugal way of studying multiple loci with a distribution of selective and dominance effects. Understanding multi-locus associations in terms of population structure arising from recent selfing history (Kelly, 1999, 2007) is a powerful but relatively under-utilized approach for studying partially selfing populations (though see (Lande and Porcher, 2015)). Extension of the ILEC (or similar) approximations to predict multi-locus associations under more complex forms of selection can provide general insight into how the interaction between population structure and polygenic selection shapes the eco-evolutionary dynamics of partially selfing populations.

## Acknowledgement

I thank Nick Barton for very useful suggestions on the model and critical comments on the manuscript.

## Supporting Information

### Identity and Linkage Equilibrium within Cohorts (ILEC) approximation

This paper introduces the Identity and Linkage Equilibrium within Cohorts (ILEC) approximation, which is a simplified version of the Inbreeding History Model (Kelly, 2007). The basic idea is to approximate the state of a large partially selfing population by neglecting correlations among the allelic states and homozygosity of different loci within each cohort of individuals who have the same *selfing age*. The selfing age of an individual is defined as the number of generations back in time to its most recent outcrossing ancestor. Thus, an individual produced by an outcrossing event in the present generation has selfing age 0, an individual produced via selfing from a parent who was itself produced by an outcrossing event in the previous generation has selfing age 1, and so on.

Individuals with higher selfing ages have a higher number of homozygous loci; this results in differences in the average fitness of cohorts of different selfing ages, if most segregating alleles are recessive. As a result, any single allele (additive or recessive) experiences effective selection that varies across cohorts, resulting in allele frequency differences between cohorts. Thus a partially selfing population can be viewed as a *structured* population consisting of cohorts with different selfing ages (or more generally different selfing histories). A structured population has non-zero linkage and identity disequilibria, even when there are *no associations* between loci within the sub-groups of the population, simply due to allele frequency or homozygosity differences between sub-groups. This is demonstrated below for the case of identity disequilibria between a pair of identical loci.

For simplicity, consider the case where individuals have only one type of locus, with one effect size. Assume that a fraction *f*_*i*_ of individuals in the population have selfing age *i*, and the average frequency of homozygous loci within the cohort with selfing age *i* is 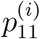. Assuming that the allelic states of loci within a cohort are uncorrelated, the fraction of individuals within cohort *i* who are homozygous at both loci for the 1 allele is 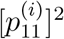. Thus, the frequency of single-locus homozygotes across the whole population is 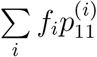, and the corresponding frequency of double homozygotes is 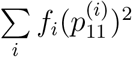. The pairwise identity disequilibrium, which is just the difference between the population-wide frequency of the double homozygote and the square of the single-locus homozygote frequency, is given by:

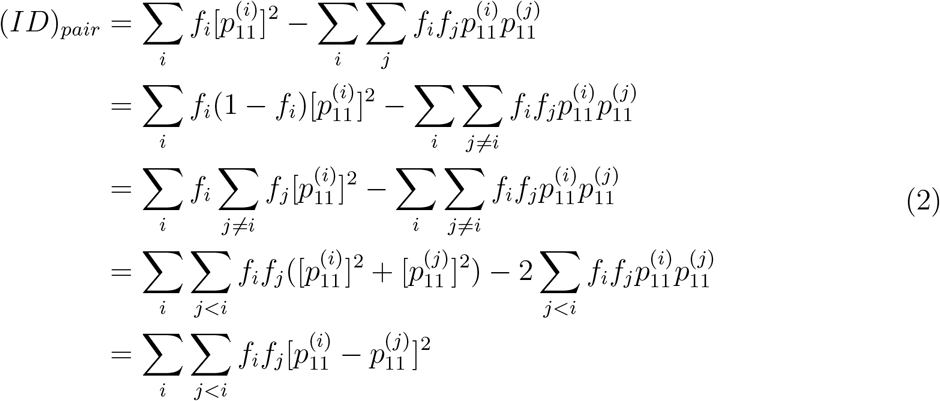

where the double summation is over all possible selfing ages. Thus, positive population-wide ID emerges, as long as the average homozygosity is different across cohorts. A similar expression for pairwise LD can also be obtained.

The assumption that the allelic states of loci within a cohort are uncorrelated is an approximation. The underlying assumption is that a single generation of outcrossing erases most associations between loci, such that the cohort of outcrossed offspring is a well-mixed group with little or no population structure. This assumption would be clearly untenable if loci are linked, or if there are epistatic interactions between loci (for example, if there were strong stabilizing selection on the additive trait). Even in the present scenario involving multiplicative selection across unlinked loci, this assumption is not strictly true, as outcrossing individuals with different selfing histories have different allele frequencies and homozygosities (resulting in differences in segregation variance among different outcrossing pairs), which can generate some structure within the cohort of outcrossed offspring. Nevertheless, this approximation generates predictions for detailed attributes of large populations, which agree quite well with results of individual-based simulations.

Under the ILEC approximation, the population is described completely by the fractions *f*_*i*_ of individuals who belong to each selfing age cohort *i*, and the frequency of heterozygous and homozygous alleles for each type of locus within each cohort. Let us denote the frequencies of homozygous and heterozygous additive loci (carrying the ‘1’ allele) within the *i*^*th*^ cohort by 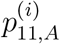 and 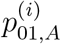, and the corresponding frequencies at partially recessive loci by 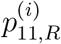 and 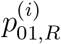. The evolution of *f*_*i*_, 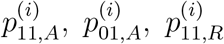 and 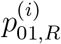 under mutation, selection and partial selfing can then be described using the following equations:

### Mutation

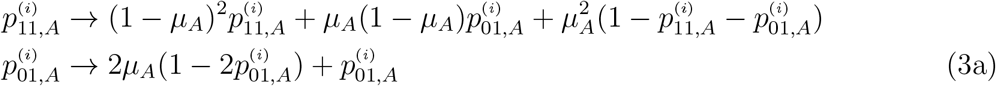

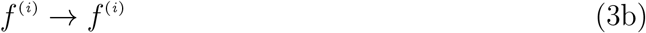

Equations (3a) represents the change in homozygote and heterozygote frequencies due to mutation at a single additive locus. A similar equation can be written for frequency changes at recessive loci, by replacing *μ*_*A*_ by *μ*_*R*_. The fraction of individuals in each cohort itself remains unchanged by mutation (eq. (3b)).

### Selection

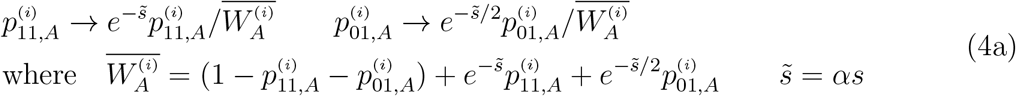

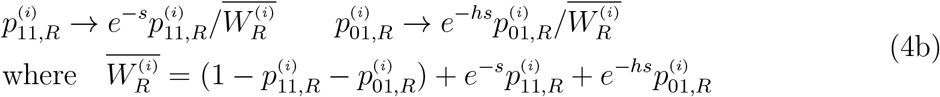

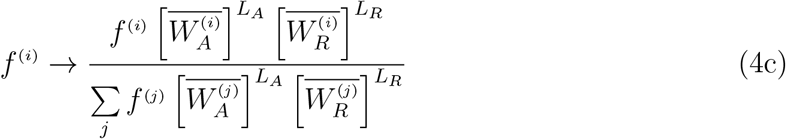

Equations (4a) and (4b) represent the effect of selection on heterozygote and homozygote frequencies at an additive locus and a recessive locus respectively, within a selfing-age cohort *i*. Equation (4c) shows how selection causes the proportions of different cohorts within the population to change in proportion to the average fitness of its members. The average fitness is determined multiplicatively by the additive and recessive loci— the fitness contribution of a single additive locus in the *i*^*th*^ cohort is denoted by 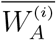 the contribution of all additive loci is 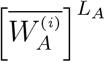, and similarly for recessive loci.

### Mating

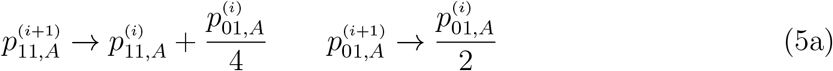

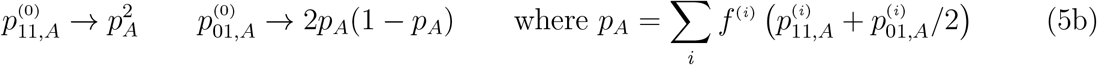

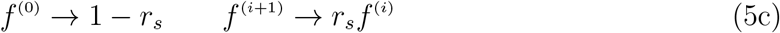

Equation (5a) shows how homozygote and heterozygote frequencies at an additive locus change due to selfing: note that the new frequencies within the (*i* + 1)^*th*^ cohort depend on the old frequencies within the *i*^*th*^ cohort. Equation (5b) shows the new heterozygote and homozygote frequencies within the outcrossing cohort: these depend only on the population-wide allele frequency *p*_*A*_, and not on the frequencies of the additive allele within each cohort separately. Equations identical to (5a) and (5b) can be written down for frequency changes at recessive loci. Equation (5c) shows the proportion of individuals belonging to different selfing age cohorts after mating; the fraction of outcrossed individuals (with selfing age 0) is just 1 − *r*_*s*_.

Equations (3)-(5) can be iterated over generations, until the proportions *f*_*i*_, and the ho-mozygote and heterozygote allele frequencies 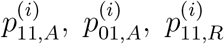 and 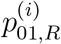 reach stationary (equilibrium) values. Note that the concept of equilibrium or steady state is not strictly well-defined for this model: starting with a fully outcrossed population (i.e., *f*_0_ = 1, *f*_*i*_ = 0 for all *i* > 0) at *t* = 0, cohorts with higher and higher selfing ages are generated in each generation. Thus, in principle, *f*_*i*_ can be non-zero for all cohorts *i* with selfing ages 0, 1, · · · *t* in generation *t*. However, the proportions *f*_*i*_ become vanishingly small for *i* ≫ −1/ log(*r*_*s*_) even in the absence of selection, while *f*_*i*_ show a steeper decline with *i* when there is selection against partially recessive alleles. Thus, in practice, all cohorts reach equilibrium if the above recursions are iterated sufficiently long. The fact that only the first few cohorts have non-zero occupancy makes this a relatively economical way of approximating population structure in a highly polygenic context.

Under the ILEC approximation, we can express the frequency of any multi-locus genotype in terms of the proportions *f*_*i*_ and the homozygote and heterozygote frequencies in each cohort. The probability *P* (*m*_11,*A*_, *m*_01,*A*_, *m*_11,*R*_, *m*_01,*R*_), that an individual has *m*_11,*A*_ homozygous and *m*_01,*A*_ heterozygous loci for the additive ‘1’ allele, and *m*_11,*R*_ and *m*_01,*R*_ homozygous and heterozygous loci respectively for the partially recessive allele, is given by:

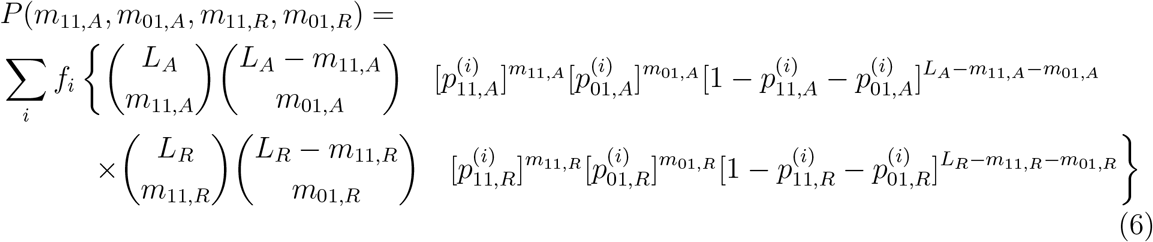

where the sum is over all selfing ages *i* for which *f*_*i*_ is non-zero. The equation above simply reflects the ILEC assumption that the states of loci within any selfing age cohort are statistically independent: then the numbers of loci (of a particular type) with states ‘00’, ‘01’ and ‘11’ must have a trinomial distribution across individuals within a particular selfing age. Equation 6 allows us to calculate any pairwise associations, as in eq. (2) above. We can also use eq. (6) to generate founder genotypes when simulating colonisation from a source population. Approximate distributions of genetic load in the population under the ILEC approximation (shown by lines in figs. 2c and 2d in the main text) were obtained by first sampling a large numbers of genotypes according to eq. (6) and then plotting the distribution of load among these.

